# Probing the Glymphatic System Using Optical Imaging and Mathematical Modeling

**DOI:** 10.64898/2026.01.22.701195

**Authors:** Micah Hesketh, Haijun Xiao, Binita Shrestha, Tyrone Porter, Carrington Lea, Binod Rizal, Peter Hinow, Muna Aryal

## Abstract

The glymphatic system is a macroscopic waste clearance pathway in the central nervous system (CNS), crucial to maintaining neural homeostasis through the exchange of cerebrospinal fluid (CSF) and interstitial fluid (ISF). Impairments in this system have been associated with neurodegenerative diseases such as Alzheimer’s and Parkinson’s, emphasizing the need to understand and potentially improve glymphatic clearance. We investigate tracer diffusion and transport in the glymphatic system by combining optical imaging in rats and mathematical modeling using a partial differential equation of reaction-advection-diffusion type. The experimental conditions differ in the levels of isoflurane-induced anesthesia (1.5, 2 and 3%) and the molecular weight of the tracer compounds (1 and 160 kDa). The optimal parameters show a close clustering of the diffusion coefficients and a wider spread of the flow velocities of the cerebrospinal fluid. Our work contributes to a better understanding of transport processes in the brain parenchyma during sleep.

## 1 Introduction

The glymphatic system is a recently discovered macroscopic waste clearance pathway in the central nervous system (CNS) (Iliff et al., 2012; Benveniste et al., 2019b). It plays a crucial role in maintaining neural homeostasis by facilitating the removal of metabolic waste through the exchange of cerebrospinal fluid (CSF) and interstitial fluid (ISF). This system functions through a network of perivascular channels: CSF enters the brain along the periarterial spaces, exchanges with ISF within the brain parenchyma, and exits along the perivenous pathways. The movement of CSF through these pathways is driven primarily by arterial pulsation, respiration, and CSF pressure gradients (Stoodley et al., 1997; Hadaczek et al., 2006; Iliff et al., 2013). A key component in this process are the aquaporin-4 (AQP4) water channels, predominantly expressed on the endfeet of astrocytes that surround the cerebral vasculature (Smith et al., 2017; Mestre et al., 2018; Mader and Brimberg, 2019; Simon et al., 2022). These channels facilitate the convective influx of CSF into the brain parenchyma, promoting the clearance of interstitial solutes such as amyloid-*β* and tau proteins. In particular, the efficiency of the glymphatic system is enhanced during sleep (Xie et al., 2013; Mendelsohn and Larrick, 2013; Benveniste et al., 2019a; Fultz et al., 2019). This is attributed to the expansion of the extracellular space, which increases the level of CSF-ISF exchange.

Dysfunction of the glymphatic system has been implicated in several neurological disorders, including Alzheimer’s disease (Kyrtsos and Baras, 2015; Peng et al., 2016; Nedergaard and Goldman, 2020; Reeves et al., 2020; Simon et al., 2022; Braun et al., 2023), Parkinson’s disease (Nepozitek et al., 2025), and traumatic brain injury (Iliff et al., 2014). Impaired glymphatic clearance can lead to the accumulation of neurotoxic waste products, which contributes to disease progression. Understanding the mechanisms regulating glymphatic function is critical to developing therapeutic strategies that aim to enhance brain waste clearance, for example, the application of transcranial ultrasound (Aryal et al., 2022; Xiao et al., 2025), electrical stimulation (Lim et al., 2023), photobiomodulation (Salehpour et al., 2022), and lifestyle choices (Reddy and van der Werf, 2020).

Mathematical modeling and simulation have long proved their usefulness in studying the complex dynamics of substance movement within the brain (Mardal et al., 2022; Bohr et al., 2022; Kelley and Thomas, 2023). A commonly used tool is the continuous approach based on partial differential equations structured by space and time. The concentrations of various chemical species change in response to convective flows (advection), dispersion or diffusion, and clearance or chemical reactions between species (Kyrtsos and Baras, 2015; Ratner et al., 2017; Ray et al., 2019; Poulain et al., 2023). Here we aid in the interpretation of the optical imagSing experimental results by constructing a reaction-advection-diffusion equation. Diffusion is the passive movement of molecules from areas of higher concentration to lower concentration due to thermal motion (Einstein, 1905) or eddies and turbulent flows (Seinfeld and Pandis, 2016). It plays a key role in the spread of small solutes over short distances within the interstitial spaces of the brain. However, diffusion alone is inefficient for transporting larger molecules or facilitating long-range clearance. In contrast, advection refers to the bulk movement of fluid that carries solutes with it. In the glymphatic system, advection and diffusion are driven by arterial pulsation, respiration, and other physiological forces. Although diffusion dominates for small molecules at microscopic scales, advection becomes increasingly important for larger solutes and transport over greater distances (Ray et al., 2019).

In this study, we employ optical imaging as a practical and versatile alternative to magnetic resonance imaging to investigate glymphatic transport. Although magnetic resonance imaging has played a key role in characterizing large-scale brain diffusion processes, its application is often constrained by high costs, technical complexity, and limited accessibility. In contrast, optical imaging offers a cost-effective and flexible platform, making it particularly advantageous for studies operating under experimental or resource limitations. Our approach enables the extraction of rich spatitemporal information from such constrained settings.

## 2 Materials and Methods

As described in (Xiao et al., 2025), this study was carried out in rats under three specific isoflurane-induced anesthetic conditions: 1.5, 2 and 3%. Throughout the study, we administered two imaging tracers, IR800 (Free IRDye800, 1 kDa) and IR800-IgG (IRDye800 labeled IgG antibody, 160 kDa) by intrathecal injection to get access to the glymphatic space. The physiological states were monitored until 70 minutes after the injection of the tracer and the distribution of the tracers in *ex vivo* brain slices was subsequently assessed and integrated with the mathematical model.

### 2.1 Delivery Agents

The near-infrared region dye IRDye^®^ 800CW NHS Ester (IR800 1 kDa) and IR800 conjugated human immunoglobulin G (IgG) antibody (IR800-IgG 160 kDa) were used as model drugs. IR800 was purchased from LI-COR Biosciences (Lincoln, NE). The conjugation of IR800 with IgG was performed by our collaborator at the University of Texas at Austin following the manSufacturer’s protocol. Human IgG was purchased from Sigma (Sigma-Aldrich, St Louis, MO). The IR800 dye was then dissolved in DMSO (10 mg/mL). Human IgG (5 mg/mL) was prepared in sodium phosphate azide-free buffer (pH 8.5). The dye was added to the IgG solution in a final ratio of 2:1. The mixture was allowed to react for two hours at room temperature in the dark. The labeled IgG was purified using a Zeba Spin^®^ desalting column 40K (Thermo Scientific, Waltham, MA). The purified labeled IgG was collected, shipped on dry ice to the Loyola University Chicago Laboratory, and stored at -20 °C until use. The absorbance of the labeled dye was measured using a UV-Vis-NIR spectrophotometer (Shimadzu, Kyoto, Japan). The degree of labeling was calculated using the following formula

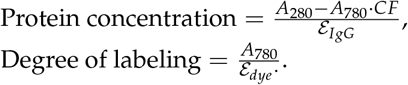

Here *A*_280_ and *A*_780_ are the absorbances of dye and IgG, respectively, and *ε*_*IgG*_ = 210, 000 M^−1^ cm^−1^ and *ε*_*dye*_ = 240, 000 M^−1^ cm^−1^ are the molar extinction coefficients for IgG and dye, respectively, in PBS. CF is the correction factor of 3% recommended by the manufacturer.

### 2.2 Characterization of Delivery Agent (IR800-IgG)

UV-Vis-NIR spectrophotometer and Sodium dodecyl sulfate-polyacrylamide (SDS-PAGE) gel electrophoresis were used to confirm the conjugation of IR800 to IgG. For SDS-PAGE gel electrophoresis, 10 *µ*g of Control IgG (unlabeled) and IR800-IgG (labeled) were mixed with 4X Laemmli sample buffer containing 0.05% volume *β*-mercaptoethanol. The mixture was heated at 90 °C for 5 minutes on a heat block. The samples were loaded on 4-20% Mini-PROTEAN^®^ TGX precast protein gels (Bio-Rad) and electrophoresis was performed at 100 mV for 75 minutes. The gels were stained with Coomassie Blue for two hours and then de-stained overnight. The gels were imaged using GelDoc^TM^ EZ imager (Bio-Rad). To confirm the fluorescence signal from IR800-IgG, the gel was imaged using an Odyssey LICOR infrared imager using an 800 nm channel. Before animal use, the stock solutions of IR800 and IR800-IgG (1 mg/mL) were prepared by individually dissolving them in DMSO, which were then diluted with artificial cerebral spinal fluid (CSF) to obtain a working solution (0.02 mg/mL). To compare the ultrasonic glymphatic-based diffusion, both IR800 and IR800-IgG were administered intrathecally according to the experimental groups. The dose of each delivery agent was 4 *µ*g/kg, and the injected volume was 80 *µ*L. Figure 1 A shows that IRDye^®^ 800CW NHS ester was conjugated to IgG through primary and secondary amines. The UV-Vis -NIR spectra of IR800-IgG demonstrate an IgG absorbance peak at 280 nm and a dye absorbance peak at 780 nm in Figure 1 B. Based on these absorbance values, the degree of labeling for IR800-IgG was found to be 2.49 moles per mole of IgG. The conjugation was further confirmed by SDS-PAGE gel electrophoresis, shown in Figure 1 C. The *β*-mercaptoethanol reduced IgG samples, both control and labeled, resulting in two bands, one from heavy chains ≈50 kDa and another from light chains ≈25 kDa. The fluorescence images of the gel show the IR800 signal in both bands, indicating that the IR800 dyes were conjugated to both heavy and light chains.

**Figure 1.**
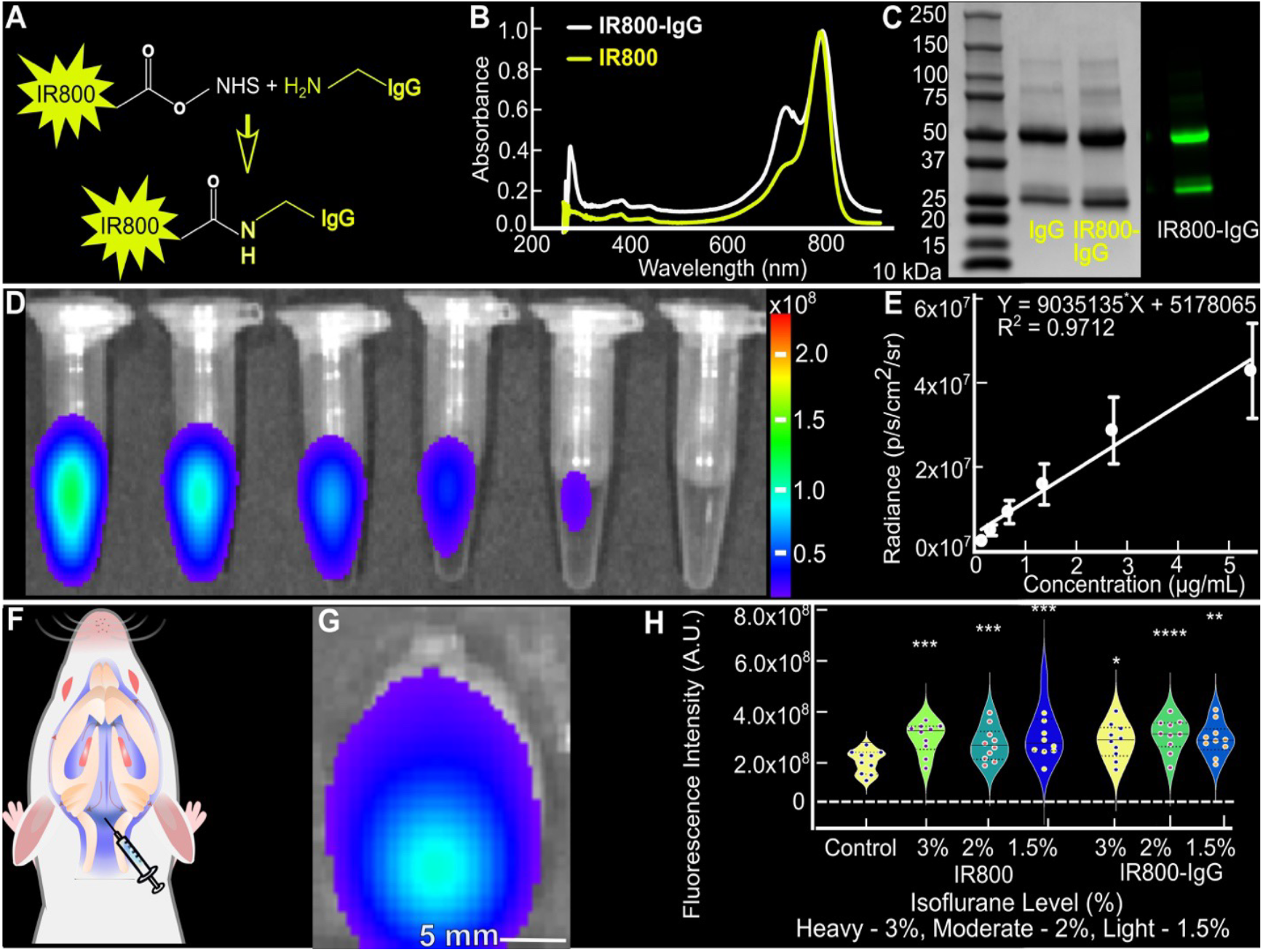
Characterization and calibration of IR800 in *in vitro* and *in vivo* settings for glymphatic transport analysis. **A** Scheme illustrating IR800 dye conjugation to IgG via NHS chemistry. **B** UV-Vis spectra of IR800 and IR800-IgG. **C** SDS-PAGE gel images showing Coomassie blue staining (left) and fluorescence at 800 nm (right). **D-E** Calibration curve for IR800 in *in vitro* settings using IVIS. **F** Schematic depiction of intrathecal injection to access imaging tracers within the glymphatic system. **G** *Ex vivo* IVIS image of a rat brain demonstrating successful IR800 delivery via intrathecal injection. **H** Calibration curve for IR800 in *ex vivo* settings using IVIS, utilizing the *ex vivo* image from (G) under varying anesthetic conditions. Data represented as mean ± SD. * *p* ≤ 0.05, ** *p* ≤ 0.01, *** *p* ≤ 0.001, **** *p* ≤ 0.0001.

### 2.3 Animals

Healthy male Sprague-Dawley rats were obtained from Charles River and used after acclimatization for three days. A total of seven experimental groups were included, one control group (no injection, serving as baseline calibration), and three groups for each tracer type (IR800 and IR800-IgG), with each group consisting of 4-5 rats, resulting in a total of 35 rats used in the study. A total of 35 subjects were initially included in the study. Subjects were excluded based on predefined criteria, including unsuccessful intrathecal injections and adverse effects associated with over-injection, such as brain swelling. The remaining study groups contributed varying numbers of rat observations, with some groups providing data from four rats and others from five rats. After applying the predefined inclusion and exclusion criteria, a total of 24 rats were included in the final analysis. The rats had an average weight of 273 ± 25 grams during the study. The use of animals in this research was approved by the Institutional Animal Care and Use Committee (IACUC) of Loyola University, Chicago, IL. Rats were divided into three main groups based on anesthesia levels: heavy (anesthesia with 3% isoflurane), medium (anesthesia with 2% isoflurane) and light (anesthesia with 1.5% isoflurane). Two imaging tracers of different sizes (IR800 - 1kDa and IR800-labeled IgG antibody-160 kDa) were injected into the spinal fluid via the cisterna magna as described below. During the experiment, careful observation of several crucial physiological indicators in rats was performed, including heart rate, respiratory rate, oxygen levels, perfusion, and body temperature. Seventy minutes after the cisternal injection, the rats were euthanized, and their brains were extracted for *ex vivo* optical imaging using IVIS. This process aimed to measure the amount of tracer delivered to the brain tissue.

### 2.4 Intrathecal Cisterna Magna Injection

The animals were placed in a chamber where they received ambient air containing 4% isoflurane through an electronic vaporizer (SomnoFlo, Kent Scientific Corporation) for around 5 minutes. Once the absence of paw reflex was confirmed, the animals were weighed and secured onto a custom-made stereotaxic frame created using a 3D printer (Ender-3, Creality 3D Technology) within the laboratory. The administration of isoflurane for the animal experiments was calibrated to meet the specified target levels using a calibrated vaporizer. To maintain a consistent level of anesthesia throughout the procedure, the flow rate of isoflurane was adjusted on the basis of the animals’ body weight. A heating pad connected to a physiological monitoring system (PhysioSuite, Kent Scientific Corporation) was placed beneath the rats’ bodies to ensure warmth and comfort. Eye lube (OptixCare Eye Lube Plus) was applied to provide lubrication and moisture. To access the cisterna magna for intrathecal injection, hair around a 1 cm area of the rat’s neck was removed using an electric hair trimmer (Philips Norelco 5500 Series), followed by the application of Nair Baby Oil Hair Remover Lotion to ensure complete depilation. The specified amount of model drug solutions was then mixed with artificial cerebral spinal fluid (Tocris, Minneapolis, MN) and introduced into the cisterna magna of the rats using a 27-gauge 13 mm butterfly needle, connected to a 12 cm polyurethane tubing (SAI Infusion Technologies, Lake Villa, Illinois) and a one-milliliter syringe via a 27-gauge 13 mm needle (PrecisionGlide, BD) as noted in Figure 1. The tracer was injected slowly over 30 s into the cisterna magna, with a total volume of 80 *µ*L, resulting in an injection rate of 2.66 *µ*L s^−1^ in order to minimize disturbance of the natural cerebrospinal fluid dynamics and avoid alterations in intracranial pressure. This rate is slightly higher than the low infusion regime (<0.12 mL min^−1^, ∼ 2 *µ*L s^−1^) reported in prior studies (Belov et al., 2021), where intracranial pressure (ICP) can be effectively regulated through pressure-dependent CSF drainage and craniospinal compliance mechanisms. In contrast, faster or bolus-like injections are known to produce transient, rate-dependent increases in ICP due to the limited capacity of the craniospinal system to immediately accommodate rapid fluid addition. However, these pressure elevations are typically short-lived and physiologically tolerated when injection volume is controlled, as compensatory mechanisms - including accelerated CSF outflow and contributions from craniospinal compliance - act to buffer the transient increase in pressure (Pass et al., 2018). Supporting this, experimental observations have demonstrated transient vascular compression lasting only a few minutes following intrathecal bolus administration, indicating rapid physiological adaptation to restore intracranial equilibrium. Therefore, although the injection rate used here is higher than slow infusion conditions, it remains within a range commonly employed in preclinical studies and is not expected to induce sustained or detrimental alterations in intracranial pressure. During the experiments, the body temperature, oxygen saturation, heart rate, and perfusion of the animals were monitored, with no significant changes observed. However, the respiratory rate differed significantly between 1.5% and 3% isoflurane. As expected, 1.5% represents a lighter level of anesthesia, while 3% corresponds to a deeper sleep-like state. Anesthesia levels also significantly affect glymphatic transport, since glymphatic flow varies with physiological state - awake, asleep, or anesthetized - and depends on the type of anesthesia used e.g. isoflurane, ketamine, propofol (Gakuba et al., 2018; Hablitz et al., 2019; Benveniste et al., 2017, 2019a; Dong et al., 2024; Wen et al., 2024).

### 2.5 Fluorescence Imaging

The extracted brains were stored in formalin for at least 24 hours before evenly sectioned into 10 slices of 2 mm thickness for the fluorescence assay. A noninvasive IVIS Spectrum Imaging System (Perkin Elmer, Waltham, MA) was used to record the total amount and distribution of dye molecules delivered to the cisterna magna based on their photon flux. The fluorescence exposure time was set to one second. The F-stop was set at two and the binning factor was four. The subject’s height was set at 0.2 cm. A pair of indocyanine green background, excitation passband: 665-695 nm, and emission passband: 810-875 nm filters were used to filter the fluorescence of IR800 dye molecules. To obtain the calibration curve for IR800, a stock soluStion was prepared by dissolving 2.0 mg of IR800 in 2.0 mL of DMSO, which was then serially diluted with artificial cerebral spinal fluid to obtain solutions with dye concentrations ranging from 20.0 *µ* g / mL to 0.16 *µ* g / mL, as noted in Figure 1 D. *In vivo* calibration of IR800 was also performed upon intrathecal injection of different sized IR800 tracers (free IR800 i.e., 1kDa, and IR800-labeled IgG antibody i.e., 160 kDa) at different anesthetic states as shown in Figure 1 G-H. The fluorescence signal was captured using the IVIS Spectrum Imaging System with the same settings described above. The total photon counts for all animals are shown in Figure 3, showing that the brain sizes and the injected amounts of IR800/IR800-IgG are comparable across all 24 animals.

### 2.6 Mathematical Model Construction and Implementation

The mathematical model is defined in a piriform domain Ω symmetric about the *x*-axis. The boundary curve is given by

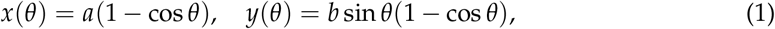

where 2*a* = 20 and 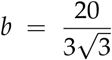 to achieve a length of 20 mm and a width of 10 mm. The orientation is so that *x* = 0 corresponds to the rostral olfactory bulb and *x* = 20 corresponds to the caudal cerebellum, see Figures 2, 4 and 7.

**Figure 2.**
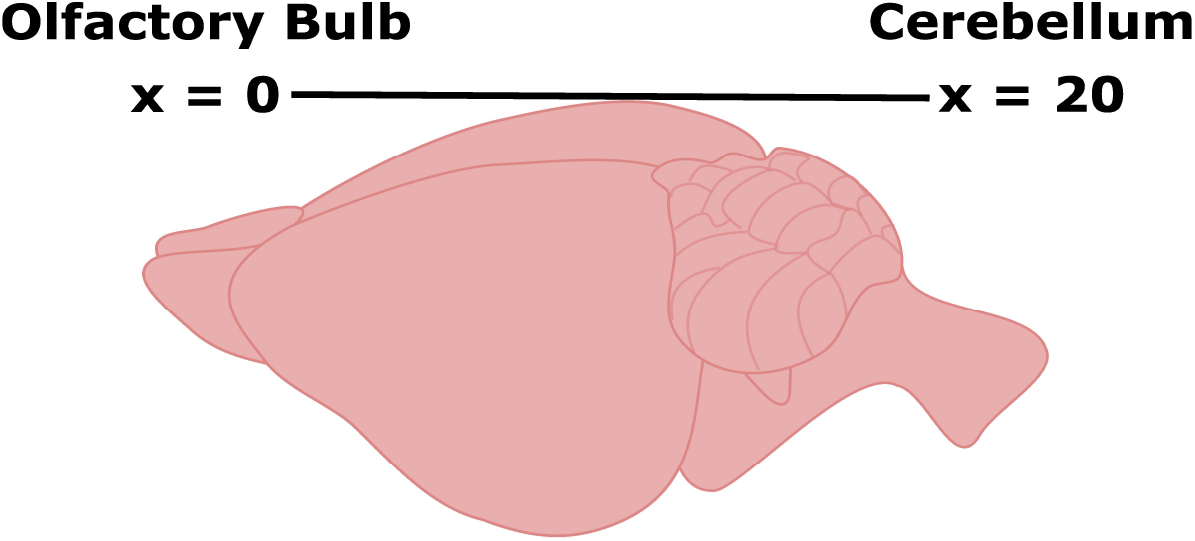
Schematic representation of an adult rat brain spanning 20 mm from the olfactory bulb to the cerebellum. Fluorescence data from 2 mm-thick *ex vivo* brain slices were used to calibrate the advection and diffusion parameters of the mathematical model.

**Figure 3.**
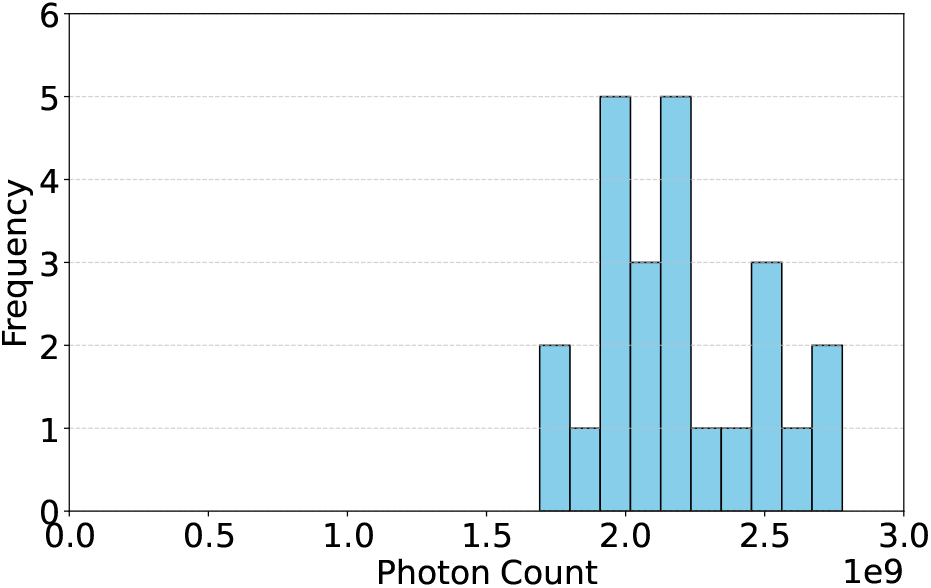
The total photon counts for 24 animals. The average is 2.2 ·10^9^ and the standard deviation is 2.9 ·10^8^, giving a coefficient of variation of 13%.

**Figure 4.**
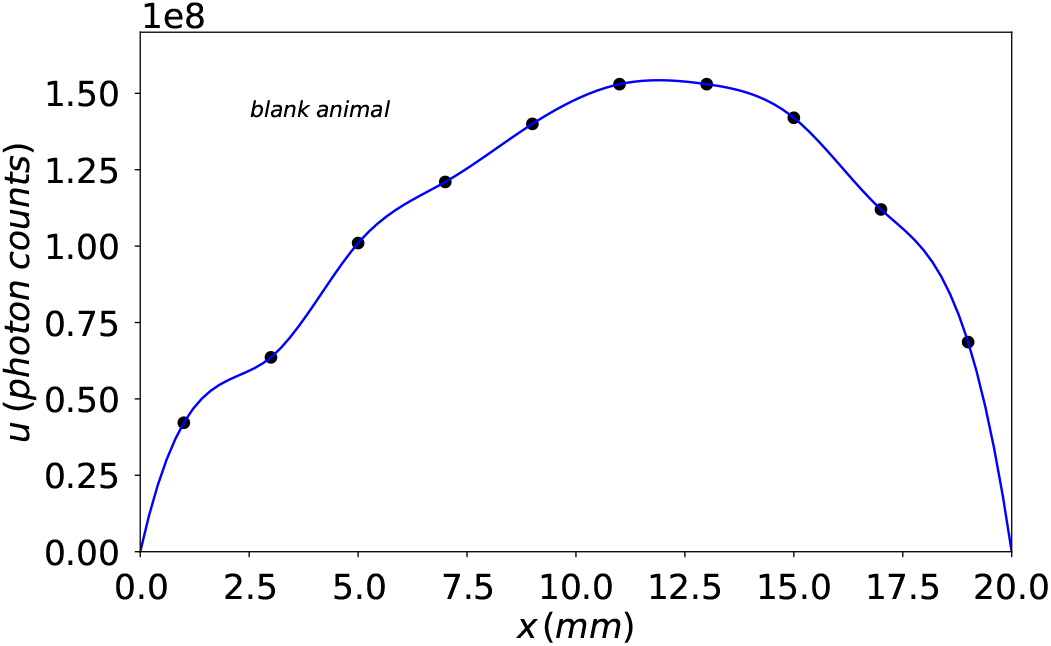
The fluorescence distribution of one blank animal fitted with a cubic spline. This supports the choice of the shape of the domain Ω in Equation (1).

The reaction-advection-diffusion equation for the concentration of a single substance *u*(*x, y, t*) is

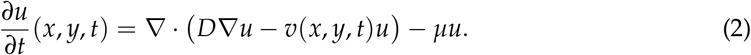

Here, *D* is the diffusion coefficient and *µ* is the clearance rate. The velocity field *v*(*x, y, t*) is selected as follows

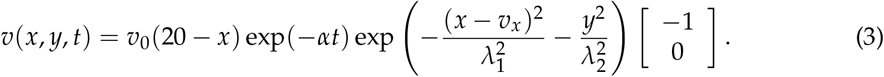

The velocity field is directed from the back to the front of the skull and is modulated by the parameter *v*_0_ (in mm s^−1^). Furthermore, it is time-dependent to account for a declining impact of the pressure increase after the tracer injection. The parameters *λ*_1_ and *λ*_2_ describe the width of the field; see Figure 7, bottom right panel. It satisfies *v*(*x, y, t*) = 0 on the boundary *∂*Ω at all times. As there is no flux of the substance across the boundary of the domain, we select homogeneous Neumann boundary conditions,

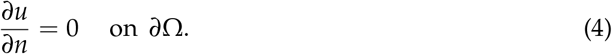

We specify the initial condition *u*(*x, y*, 0) = *u*_0_(*x, y*) by

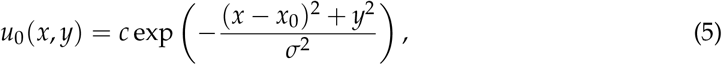

where *c* is a constant chosen to match the evolution of the total mass that varies from animal to animal. The localization and parameter *x*_0_ is chosen as part of the fitting procedure outlined below, subject to the constraint that the injection of IR800 and IR800-IgG takes place into the cisterna magna towards the back of the rat’s skull; see Figure 1 F. The shape parameter *σ* is set to 1 mm throughout. Regarding the clearance rate, (Plá et al., 2022) report a half-time of fast kinetics *T*_1/2_ = 50 min for Direct Blue 53 (molecular mass 961 Da). From this we obtain

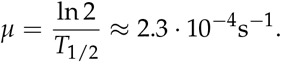

Assuming that there is no other loss or gain in the domain Ω or beyond its boundary, we have that the total mass

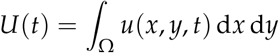

evolves according to

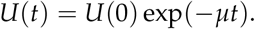

From this we find the value *c* in Equation (5) by taking the value *U*(*t* = 4200 s) as the terminal condition. With the choice of *µ* as above, we have that *U*(0) is approximately 2.63 times the terminal value.

The mathematical model from Equations (2)-(5) is solved using the Finite Volume Method (FVM) implemented in the FiPy package (Guyer et al., 2009). The coding process was considerably sped up using Gemini 3.1 Pro (Gemini Team, 2023). All python code was carefully checked. The default time integration scheme of the FVM is the backward Euler method. The domain Ω is partitioned into 10 slices of equal width orthogonal to the *x*-axis and the model soutput is integrated over each slice. Fitting the model output at *t* = 4200 s to the observations by minimizing the *L*^2^-error is done with the Nelder-Mead algorithm implemented in scipy (Press et al., 2007; Virtanen et al., 2020). The fitting parameters are *D, v*_0_, *x*_0_, *λ*_1_, *λ*_2_, and *α*, while the localization parameter of the velocity field is set to *v*_*x*_ = *x*_0_ −1. The raw data, python code and the full set of fitted parameters will be available on request and on figshare.com/s/eb7e87380e319ef9979a.

## 3 Results

Figure 4 shows the background fluorescence of a blank animal, supporting our choice of the piriform shape of the computational domain. Figure 5 shows sample images of brain slices indicating an anatomical and morphological coherence of the brains across the animals.

**Figure 5.**
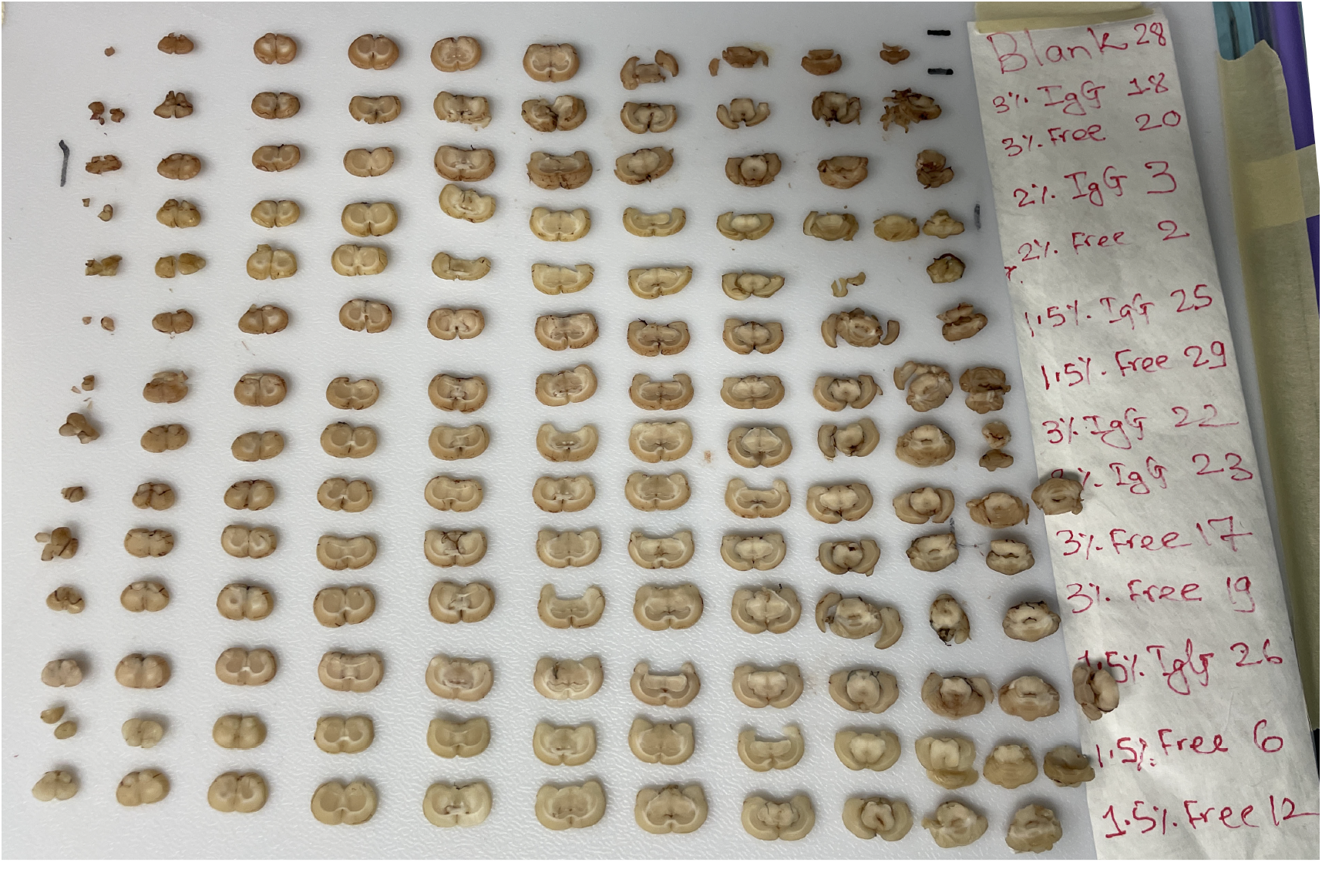
Sample images of brain slices.

Figure 6 shows the complete fluorescence distributions at the time of extraction *t* = 70 min after injection. These pictures show a high degree of coherence between multiple individual animals. Visible is a general pattern where the peak of the distribution is between 12 and 15 mm in the interior of the domain. An exception is the case of IR800 under light anesthesia where all peaks are located at the right boundary.

**Figure 6.**
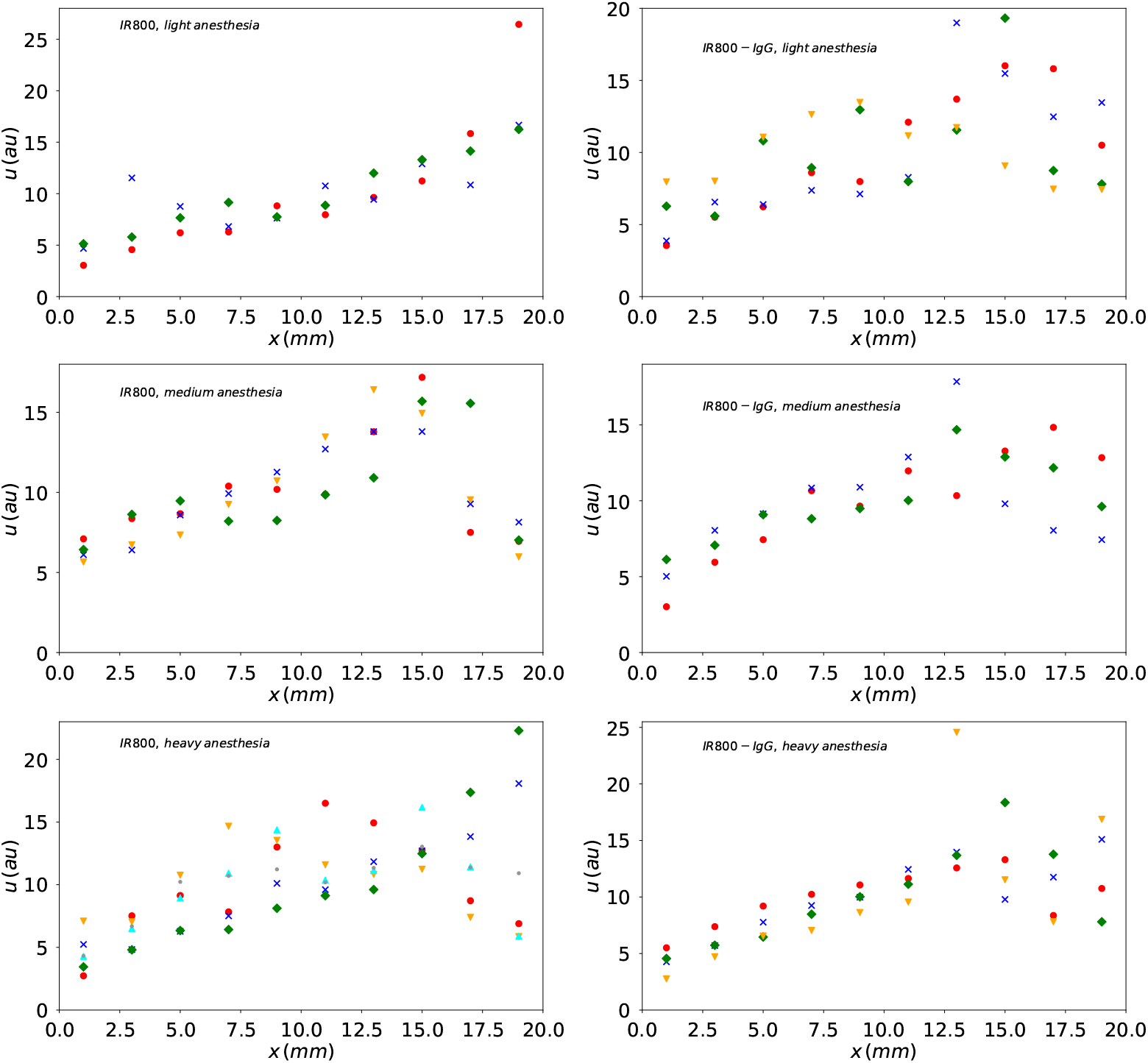
The complete fluorescence distributions at *t* = 70 min normalized to total mass 100. Each set of symbols corresponds to one animal.

Figure 7 shows a sample simulation together with the normalized fluorescence data of animal 13 (IR800, heavy anesthesia, individual 6). The corresponding figures for all other animals are part of the data depository. The medians and interquartile ranges for all parameters are reported in Table 1. These are chosen as they are more robust to outliers than mean and standard deviation. Table 1 also contains the values of *D* and *v*_0_ for the subsets of animals injected with IR800 and IR800-IgG, respectively. The only difference is that the median value of *v*_0_ is some-what higher for the heavier molecule IR800-IgG than for the lighter molecule IR800. Figure 8 shows a plot of all 24 optimal (*D, v*_0_) pairs.

**Figure 7.**
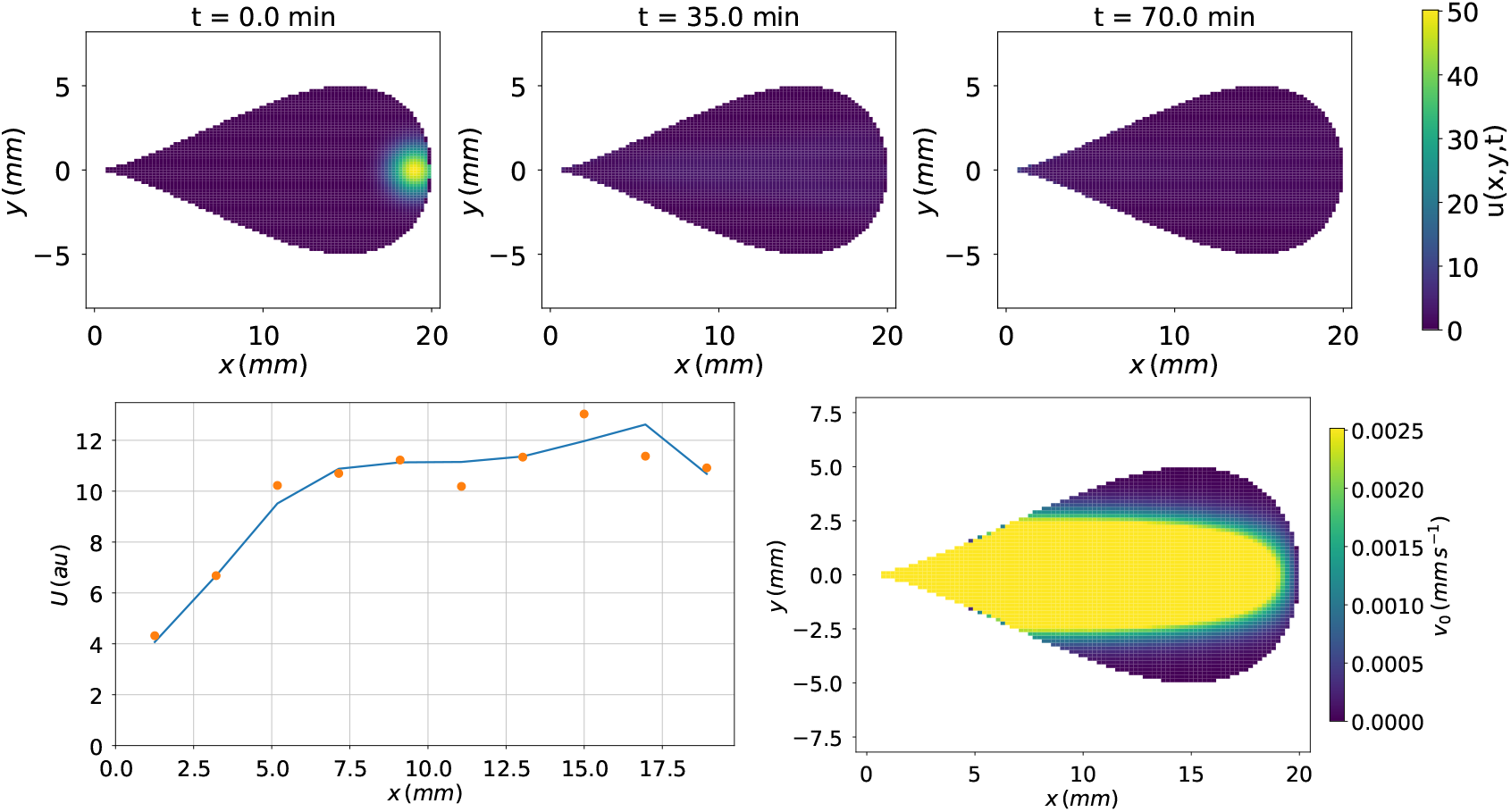
**(Top row.)** Spread of the tracer particles on a piriform shape with length 20 mm and width 10 mm. The parameters are *D* = 1.3 10^−4^ mm^2^ s^−1^, *v*_0_ = 2.5 10^−3^ mm s^−1^, *x*_0_=19 mm, *λ*_1_=14.6 mm, *λ*_2_=1.76 mm and *α* = 8.8 10^−4^ s^−1^. **(Lower left.)** The integrated solution over 10 parallel slices of equal width at the terminal time (solid curve) together with the normalized fluorescence data of animal 13 (IR800, heavy anesthesia, individual 6; dots). **(Lower right.)** The magnitude of the velocity field at time *t* = 0 as described in Equation (3). The direction of the flow is from right to left and parallel to the *x*-axis.

**Table 1:** Medians and interquartile ranges for all parameters. The values of *D* and *v*_0_ are also listed for the subgroups injected with the small and the large tracer molecule.

| Parameter | Median ($Q_2$) | IQR ($Q_3 - Q_1$) |
| --- | --- | --- |
| $D$ | $1.28 \cdot 10^{-4} \text{ mm}^2 \text{ s}^{-1}$ | $1.64 \cdot 10^{-4} \text{ mm}^2 \text{ s}^{-1}$ |
| $v_0$ | $2.56 \cdot 10^{-3} \text{ mm s}^{-1}$ | $6.0 \cdot 10^{-3} \text{ mm s}^{-1}$ |
| $x_0$ | 18.75 mm | 2 mm |
| $\lambda_1$ | 16.9 mm | 4.8 mm |
| $\lambda_2$ | 1.39 mm | 1.2 mm |
| $\alpha$ | $9.9 \cdot 10^{-4} \text{ s}^{-1}$ | $2.1 \cdot 10^{-3} \text{ s}^{-1}$ |
| $D$ (IR800) | $1.29 \cdot 10^{-4} \text{ mm}^2 \text{ s}^{-1}$ | $2.0 \cdot 10^{-4} \text{ mm}^2 \text{ s}^{-1}$ |
| $D$ (IR800-IgG) | $1.22 \cdot 10^{-4} \text{ mm}^2 \text{ s}^{-1}$ | $1.11 \cdot 10^{-4} \text{ mm}^2 \text{ s}^{-1}$ |
| $v_0$ (IR800) | $2.44 \cdot 10^{-3} \text{ mm s}^{-1}$ | $3.4 \cdot 10^{-3} \text{ mm s}^{-1}$ |
| $v_0$ (IR800-IgG) | $3.59 \cdot 10^{-3} \text{ mm s}^{-1}$ | $1.0 \cdot 10^{-2} \text{ mm s}^{-1}$ |

**Figure 8.**
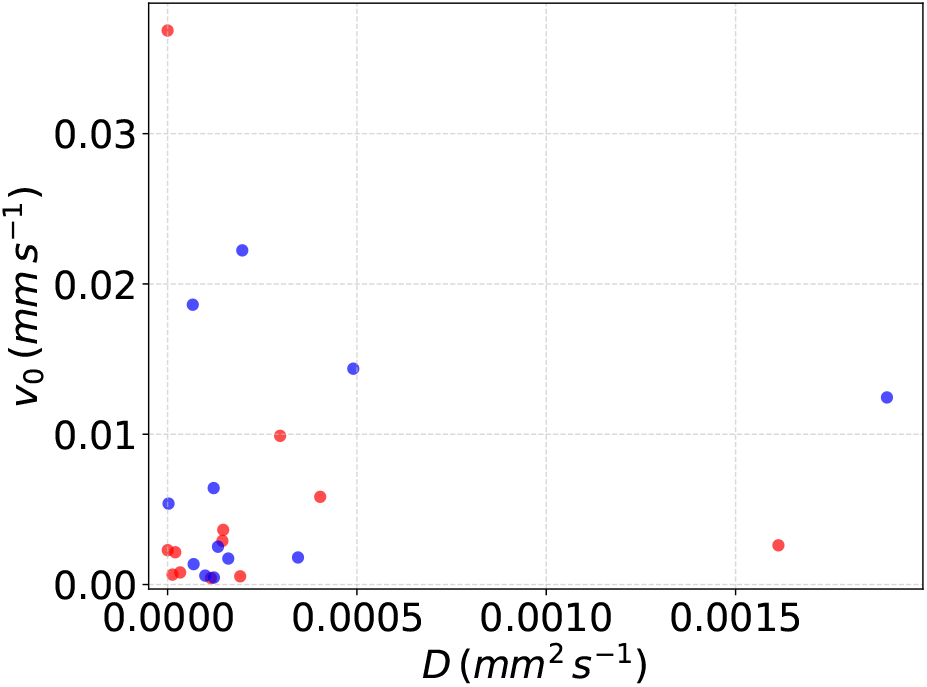
The optimal fits of *D* and *v*_0_ for all 24 animals. Values for IR800 are marked in red, values for IR800-IgG are marked in blue.

## 4 Discussion

In this study, we combine *ex vivo* fluorescent imaging and mathematical modeling to investigate the transport of solutes within the glymphatic system under varying physiological states. For two tracers of distinct molecular sizes, IR800 (1 kDa) and IR800-IgG (160 kDa), we observe no differences in the diffusion parameter *D*. The parameter of the velocity field at the initial time *v*_0_ is only slightly higher for IR800-IgG than for IR800. In earlier work, (Xie et al., 2013) found that sleep or sleep-like states expand the extracellular space, thus facilitating glymphatic transport. Future work is needed to better quantify a possible increase in porosity with increasing depth of anesthesia and with different anesthetic compounds (Hablitz et al., 2019). The dependence of glymphatic kinetics on molecular weight dis a topic of active research (Miao et al., 2024; Plá et al., 2025). However, *ex vivo* optical imaging is unsuited to address this issue with the necessary spatial and temporal precision.

Our mathematical model captures tracer dynamics over a 70-minute snapshot and uses a back-to-front time-dependent velocity field. While the two-dimensional computational domain is a simplification, it captures well the specific geometric shape of the rat skull. Our optimal values for *D* are comparable to those

1. used by (Ray et al., 2019) and (Poulain et al., 2023) in their numerical simulations, e.g. 1.03·10^−4^ mm^2^ s^−1^ for ^14^C-inulin,
2. reported by (Ray et al., 2021, Table 1) from dynamic contrast-enhanced magnetic resonance imaging experiments,
3. the diffusion coefficient of 4·10^−4^ mm^2^s^−1^ for the 480 Da Rhodamine 6G molecule (Gendron et al., 2008), and
4. the value of 1.36·10^−4^ mm^2^ s^−1^ for the 4 kDA FITC-dextran molecule measured by photobleaching in an agarose gel brain phantom (Miao et al., 2024).

This supports the validity of our experimental conditions and the modeling approach. Based on the experimental setup, our assumption is that the tracer injection is localized to a slice of about 2 mm in width in the back of the skull. After 70 minutes, the tracer particles have spread widely throughout the brain. Dispersive motion in glymphatic space is possibly the result of isotropic pulsatile pressure changes originating from the circulatory system during every systole and diastole. An analogy is the kneading of a piece of dough or mixing liquids with a pipette by repeatedly moving the plunger up and down. Physiological brain pulsations and their impact on fluid motion and transport processes are an active topic of interdisciplinary research (Kiviniemi et al., 2025; Lecchini-Visintini et al., 2025).

The influence of tracer delivery on intracranial pressure and glymphatic transport remains a topic for active research (Vinje et al., 2020; Belov et al., 2021; Plá et al., 2022). Our modeling and simulation results in the parameter *α* which indicates that a velocity field directed from the back to the front (which is not compensated by a flow in the opposite direction) fades away with a half-time of approximately 11 minutes. Although our results provide useful information about solute distribution shortly after injection, a more comprehensive understanding will require longer observation periods and potentially the integration of multimodal imaging data. Studies using advanced techniques such as magnetic resonance imaging e.g. (Ray et al., 2019), allow for finer spatial and temporal resolution, but come at a significantly higher resource cost. Our approach, on the contrary, demonstrates the potential of extracting meaningful quantitative insights from more accessible experimental setups. Naturally, future work should focus on discovering convergent patterns derived from different imaging modalities (Bohr et al., 2022). A deeper understanding of fluid and solute transport processes in the brain parenchyma will lead to significant progress in clinical applications such as drug delivery, stroke, and neurodegenerative disorders (Mestre et al., 2020; Bohr et al., 2022).

Recent advances have highlighted the potential of noninvasive techniques, such as focused ultrasound (FUS), to modulate glymphatic transport. Studies have shown that FUS, with (Meng et al., 2019; Lee et al., 2020; Wu et al., 2024; Gong et al., 2025) or without (Aryal et al., 2022; Ye et al., 2023; Xiao et al., 2025) microbubble-mediated opening of the blood-brain barSrier (BBB), can enhance glymphatic transport by mechanically stimulating perivascular spaces. SThis promotes the movement of CSF and ISF within the cerebrovascular compartments, offering a promising therapeutic avenue for neurodegenerative diseases characterized by glymphatic dysfunction.

## Acknowledgments

The authors extend their gratitude to the In Vivo Imaging Center at Stritch School of Medicine at Loyola University Chicago for generous provision of the IVIS system and access to the animal facility. We also appreciate the contributions and support of all members of the Aryal Lab, whose insightful discussions and equipment usage were invaluable. The authors thank Dr. Philip Chang (Department of Physics, UWM), Liam Jemison, and Berkant Palazoglu (Department of Mathematical Sciences graduate students, UWM) for valuable help. Financial support for this work was provided by the Engineering Department’s Faculty Start-Up fund at Loyola University Chicago, the Department of Chemical, Biological, and Bioengineering at North Carolina Agricultural and Technical State University, as well as support from the National Academy of Medicine Healthy Longevity Catalyst Award and the Focused Ultrasound Foundation Global Intern program.

## Statement Concerning the Use of AI

The coding process was considerably sped up using Gemini 3.1 Pro (Gemini Team, 2023). All `python` code was carefully checked.

